# Sphingosine-1-Phosphate Restrains Foamy Macrophage Formation and Modulates Inflammation in Multidrug-Resistant Tuberculosis

**DOI:** 10.64898/2026.10.15.759318

**Authors:** Abhishek Kumar, Niharika Sharma, Lalit Kumar Singh, Vijay Hadda, Amit Kumar Singh, Hridayesh Prakash

## Abstract

The high abundance of lipid loaded foamy macrophages in the granulomatous lesions is a key hallmark of advanced pulmonary tuberculosis. These macrophages serve as lipid-rich intracellular hiding niches that provides *Mycobacterium tuberculosis* (*M.tb*) survival and persistence during prolonged anti-tubercular treatment. Persistent infection and dysregulated inflammatory responses further complicate disease control, particularly in multidrug-resistant tuberculosis (MDR-TB), highlighting the need for host-directed therapeutic strategies. Our previous preclinical study demonstrated the therapeutic potential of the host sphingolipid sphingosine-1-phosphate (S1P) in reducing the mycobacterial burden in *M.tb*-infected animals, accompanied by enhanced pulmonary macrophage activation. To explore its translational potential, we investigated whether S1P modulates foamy macrophage formation in CD14 monocyte-derived macrophages from healthy donors and patients with MDR-TB. Notably, macrophages derived from patients with MDR-TB exhibited greater intracellular lipid droplet accumulation than those derived from healthy donors. This phenotype was further characterized by increased nitric oxide production and quantitative RT-PCR analysis of genes associated with immune regulation and lipid metabolism, including *APOE, LDLR1, CLEC4E, CLEC7A, GP96, IL4*, and *IL13*, which suggested a Th2-associated immunological bias in foamy macrophages. Collectively, these findings highlight the potential role of host sphingolipids in modulating foamy macrophage phenotypes and associated immunometabolic dysregulation, supporting further investigation of S1P as a host-directed therapeutic strategy against MDR-TB.

## Introduction

Emergence of drug-resistant *Mycobacterium tuberculosis* (*M.tb*) cases over last few years has become a most serious threat to the public health. Drug resistance tuberculosis burden ranges between 6.8% to 18% of tuberculosis (TB) cases worldwide. Current treatment regimens involve long treatment duration with multiple drugs, and bacteria frequently develop several mechanisms for securing their opportunistic survival during antibiotics treatment (1, 2). Drug resistant *Mycobacterium tuberculosis* has the capability to adapt in metabolically diverse intracellular environments, in which *M.tb* sabotage or evade host immune effector mechanisms (3, 4). *M.tb* bacilli can harbour inside different cells e.g., alveolar macrophages, interstitial macrophages, neutrophils, mesenchymal stem cells, and hepatocytes. Among these intracellular niches, foamy macrophages (FM) are primary host cell niche for *M.tb* inside TB granuloma. Foamy macrophages are lipid enriched cells that exhibits elevated lipid metabolism, uptake and storage (5, 6). *M.tb* exploits host lipid metabolic pathways to promote FM formation, which is a drug-inaccessible niche which provides essential nutrients to sustain long-term bacterial survival (7, 8). FM provides a favourable environment for pathogen proliferation and serves as a reservoir for subsequent granuloma seeding (5, 9). Foamy macrophages are plastic in nature and exhibit moderate to high Th2 bias in latent / MDR cases. Due to this, they contribute significantly to drug resistant and immune evasion mechanism of MDR-PTB (10). Therefore, interventions targeting both cellular and immunological aspects that contribute to the macrophage plasticity may serve a potent adjuvant therapy for MDR-PTB (11).

Host-directed therapy (HDT) has emerged an effective and alternative approach for augmenting the impact of anti-tubercular therapies. Unlike, antimicrobial drugs, HDT is concerned with improving host cell process (e.g., inflammation, macrophage homeostasis, cellular metabolism, and immune function) to enhance the host cell ability to control infection, potentially improving anti-tubercular therapy (12). In this regard, our pioneer work has demonstrated the therapeutic impact of metabolic reprogramming of *M.tb* infected animals by Sphingosine 1-phosphate (S1P) in animal model of TB (13). Sphingolipid signalling is one of the metabolic pathways involve in macrophage homeostasis and is a major regulator of cellular phenotype, cell death, inflammation, cellular trafficking and responses to infection (14, 15). Sphingolipids are the emerging regulators of macrophage plasticity, inflammatory signalling, and response to infection that could serve as a target for therapeutic intervention and the development of new adjunctive TB therapy. Sphingosine 1-phosphate is the most abundantly expressed sphingolipid mediator and has been shown to have potent immunomodulatory effects (16). In contrast to other sphingolipids with more specialized cellular functions, S1P works as an immune cell messenger and morphogen of cellular differentiation. Metabolic alterations in sphingolipid pathway can impact the functions of macrophages and inflammatory responses (17). Previously, we have shown sphingolipid mediated host immunomodulation as anti-tubercular host directed therapy. Following our central hypothesis, this study prudently demonstrated that treatment of foamy macrophages with S1P, potentially disrupted foamy phenotype and inhibited pathogenic inflammatory response. Thus, modulation of host-S1P axis could be a strategy to alter FM from a metabolically permissive environment to more conducive antimicrobial and effective immune response.

## Materials and Methods

All reagents including Histopaque (10771), Palmitic acid (P5585), Lipopolysaccharide (L6511), Sulphanilamide (S9251), N-(naphthyl ethylenediamine dihydrochloride) (684147), Formaldehyde (252549), Oil Red O solution (O1391), Sphingosine 1-Phosphate (S9666) and L-serine (S-4500) were purchased from Sigma-Aldrich. RPMI-1640 (SH30027001), 10% fetal bovine serum (10270-106) and 1% penicillin-streptomycin (15140-122) (Cytiva Hyclone, Marlborough, MA). Oxy LDL (L-34357) was purchased from Invitrogen, Isopropanol (67800) purchased from SRL and M-CSF (30025) procured from Peprotech. cDNA synthesis kit (1708890) and SYBR green (1725271) were purchased from Bio-Rad. RT-PCR primers for foamy macrophages were purchased from Barcode Biosciences. All consumables and plasticwares were obtained from Genaxy and Trueline.

### Study Participants

Peripheral Blood Mononuclear Cells (PBMCs) were collected from both male and female having MDR-PTB (n=5). MDR-PTB patient’s age between 18–55, were screened from Directly Observed Treatment (DOTS) centre of All India Institute of Medical Science (AIIMS), New Delhi. Inclusion criteria of these patients were having family history of TB, relapse, treatment ongoing, pulmonary TB (MDR-PTB) patients with MDR-PTB, positive sputum Acid fast bacilli (AFB) stain and chest X-Ray for MDR-PTB patients, CBNAAT (Cartridge-Based Nucleic Acid Amplification Test) positive and resistant to first line drugs i.e., isoniazid and rifampicin. Patients having chronic obstructive pulmonary disease (COPD), atypical pneumonia, aspergillosis, lung cancer and history of HIV positive were excluded in this study. Healthy donors with same age group were recruited (n=5). The study comprised members of the healthy group who were purified protein derivative (PPD) skin test negative and had no history of tuberculosis, mycobacterial infections, or other infectious disorders.

### Isolation and Culturing of Peripheral Blood Mononuclear Cells

Blood was collected into EDTA-vacutainer tubes for PBMC separation. PBMCs were isolated with the Histopaque-1077 density gradient and centrifuged for 20 minutes at 450x g at room temperature. The subsequent layer, which was highly enriched with monocytes was washed 3 times with Roswell Park Memorial Institute 1640 (RPMI 1640). Monocytes were cultured for four days at density of 1x10^5^ cells/well in the 96-well plate. To induce monocytes to transform into macrophages, cells were resuspended in complete RPMI 1640 (cRPMI 1640) which consists of 10% heat-inactivated Fetal Bovine Serum (FBS) and 1% penicillin – streptomycin and were incubated at 37° C in a humidified 5% CO_2_ incubator. After three hours of incubation cells were stimulated with macrophage colony-stimulating factor (M-CSF) 20 ng/ml, which was replenished every two days (18, 19).

### Foamy Macrophage Induction by Lipid Treatment

For the induction of foam cell formation, adhered macrophages were stimulated with different lipid molecules like oxidised LDL (oxyLDL) (5 µg/ml), lipopolysaccharide (LPS) (100 ng/ml), palmitic acid (PA) (100mM), naïve LDL (nLDL) (100 µg/ml) and oleic acid (OA) (200 µM) for 24 hours in 5% CO_2_ incubator at 37^°^ C to induce foamy macrophage phenotype and lipid droplet accumulation in macrophages (20–23). Untreated macrophages were served as experimental control. Oil red staining was done to confirm the foamy macrophage phenotype (24). Following the induction of foamy macrophages by different lipid molecules, cells were subjected to metabolic intervention by S1P. It has previously been demonstrated that S1P is the bioactive lipid mediator which affects macrophage polarization (16). Macrophages which induces foamy phenotype was treated with S1P (5 µM) for 24 hours at standard culture conditions. Effect of S1P on foamy macrophages was confirmed by Oil red staining and RT-PCR.

### Oil Red O Staining

Oil Red O staining was used to demonstrate the accumulation of lipid loaded macrophages. The working solution of Oil Red stain must be made fresh and should be used within two hours. After two PBS washes, cells were fixed in 10% formalin for 30 minutes, then rinsed twice with Milli-Q water (25, 26), and finally washed with 60% isopropanol for 5 minutes to make neutral lipid staining faster. The cells were examined under a phase contrast microscope after being treated with a working Oil Red solution for ten minutes and then washed three to four times with Milli-Q water.

### Estimation of Nitric Oxide

Supernatants were collected after 24 hours of lipid treatment for analysis of Nitric Oxide (NO). Using the Griess reagent method, NO release was determined by assessing the build-up of its stable oxidation product, nitrite (27). Griess reagent: 0.1% (w/v) N-(1-Naphthyl) ethylenediamine dihydrochloride (NEDA) solution in water, 1% (w/v) sulphanilamide solution in 5% (v/v) phosphoric acid was added to obtain Griess reagent (28). Therefore, 50 µl of samples and standards (200 µM, 100 µM, 50 µM, 25 µM, 12.5 µM, 6.25 µM and 3.125 µM) were added to each well of 96 well plate. Mix R1 (NEDA) and R2 (Sulphanilamide) in 1:1 ratio to make Griess reagent. Subsequently 50 µL of Griess reagent was added to each well. Then absorbance was measured at 540nm.

### Extraction of RNA and Real time PCR analysis

RNeasy spin kit was used to extract total RNA. One microgram of total RNA was used for cDNA synthesis using iScript cDNA Synthesis kit. Quantitative Reverse-Transcription Polymerase Chain Reaction (qRT-PCR), SsoAdvanced Universal SYBR Green Supermix, Quant Studio5 used to measure relative mRNA levels (18, 29). Primers for foamy macrophages are *APOE*, *CLEC4E*, *CLEC7A*, *GP96*, *LDLR1*, *IL-4*, *IL-13* (**Supplementary Table 1**) were designed using origene and purchased from barcode biosciences. The 2^-ΔΔ^Ct method was adopted to analyze relative gene expression; which was normalized to the expression of beta-actin for macrophages.

## Results and Discussion

### S1P tweak immunomodulatory response in MDR MDR-PTB patients

Competitive analysis of CD14+ monocytes from healthy and MDR-MDR-PTB patients revealed that macrophages from both healthy and MDR-MDR-PTB patients produced NO quite fast and robustly in response to S1P stimulation suggesting their sensitivity towards Sphingolipids in general. Their response toward LPS remained less spontaneous. Interestingly macrophages from MDR-MDR-PTB patients maintained high level of NO over healthy donors when co-stimulated with S1P and LPS. Interestingly similar stimulation inhibited NO titers in the macrophages from healthy donors. This observation indicated high grade redox signalling in the macrophages from MDR-PTB patients over healthy group **(Figure 1A, B)**. This contrasted with NO response which was quite high in these macrophages. We predicted Th2 or M2 programming of macrophages from MDR-PTB group. To substantiate this doubt, we quantified the secretion of IL-10 in the culture supernatant of these macrophages. Following our prediction, macrophages from MDR-MDR-PTB patients produced more IL-10 when stimulated with S1P which indicated M2 programming of macrophages from MDR-MDR-PTB patients **(Figure 1C, D)**. Based on Th2 programming of macrophages from MDR-MDR-PTB patients, we anticipated Th17 responses which promote pulmonary fibrosis in these patients and contribute to the antibiotic’s resistance. These responses also promote the foamy conversion of infected macrophages in MDR-MDR-PTB patients. To substantiate this, we quantified the ability of macrophages from MDR-MDR-PTB patients to secrete IL-6 which triggers pathological response in the patients by promoting mycobacterial burden and pulmonary fibrotic lesions. Following our prediction, unstimulated macrophages from MDR-MDR-PTB patients over healthy donors produced high IL-6 titre. Interestingly S1P was able to reduce over the period indicating immune conditioning potential of S1P in the macrophage from MDR-PTB patients **(Figure 1E, F)**.

**Figure 1.**
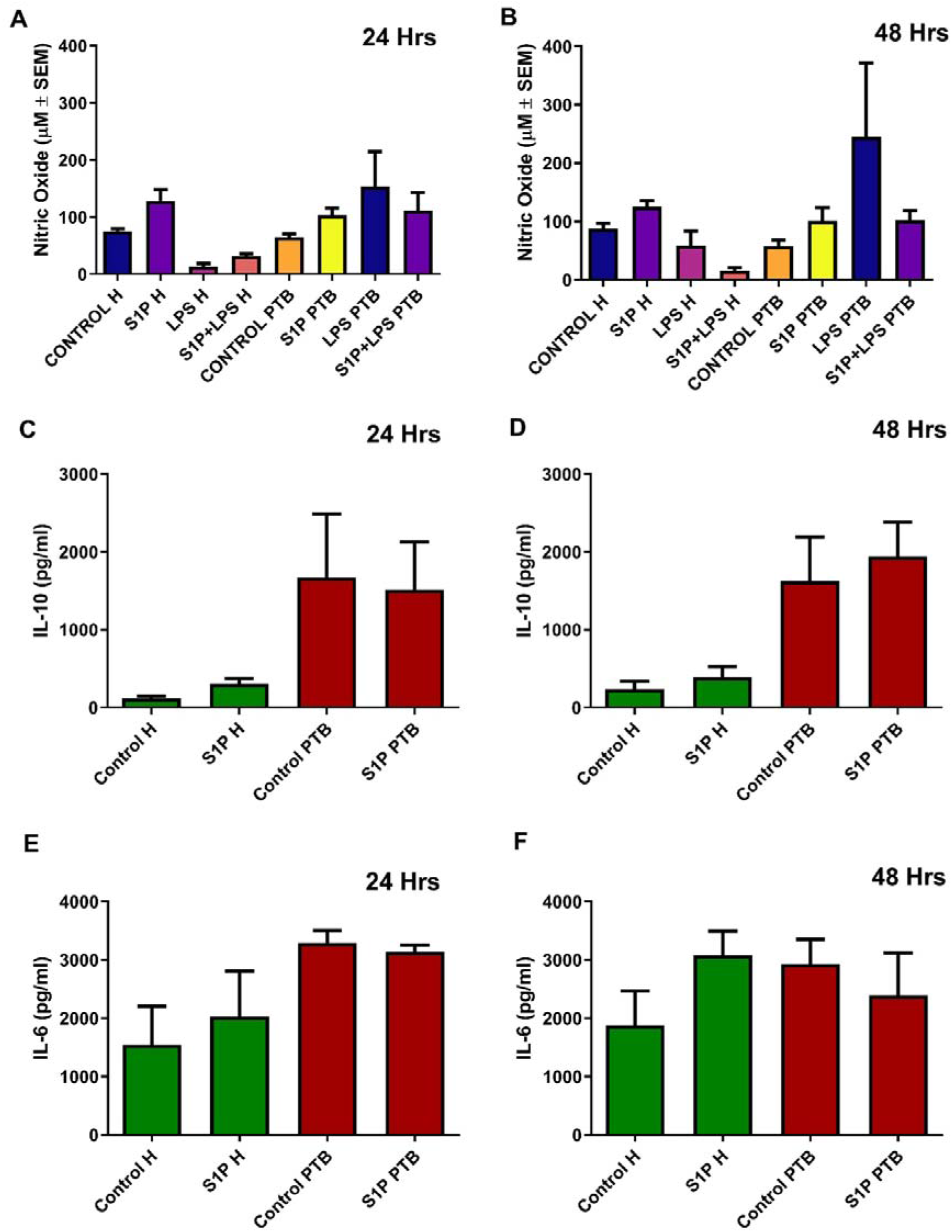
S1P treatment increases NO production and Inflammatory Cytokine levels. **(A-B)** CD14^+^ macrophages were purified from both healthy donors and MDR-PTB patients, cultured and stimulated as indicated. Their culture supernatants were collected at indicated intervals and production of NO was quantified in their culture supernatant by Griess reagent method. Shown here is the mean of µM ± SEM from indicated number of donors. **(C-D)** Healthy donors and MDR-MDR-PTB patients culture supernatants were collected at indicated intervals and secretion of IL-10 was quantified in their culture supernatant by ELISA kit from R&D system as per their protocol. Shown here is the mean of µM ± SEM from indicated number of donors. **(E-F)** Secretion of IL-6 was quantified in their culture supernatant by ELISA kit from R&D system as per their protocol. Shown here is the mean of µM ± SEM from indicated number of donors.

### S1P reduces the Lipid Accumulation and Foamy Macrophage Formation

To evaluate this, we first established the protocol of foamy differentiation from PBMCs derived monocyte from both healthy donors and MDR-MDR-PTB patients. PBMC from HD and MDR-PTB groups were cultured and treated with M-CSF. Over time, a series of morphological changes were recorded, demonstrating adhesion, spreading, and maturation into macrophages **(Supplementary Figure 1A)**. Differentiated macrophages formed distinctive internal lipid droplets in response to additional lipid-inducing stimuli, indicating the formation of foamy macrophages. Representative phase-contrast images illustrating the morphological differences between the healthy and MDR-MDR-PTB groups were observed. Phase-contrast images reveal clear morphological variations in the groups’ foam cell growth and lipid accumulation **(Supplementary Figure 1B)**. Increased foam cell production in TB patients’ macrophages indicates changed immune activation and lipid metabolism, which reflects the pathogen-driven modification of host lipid pathways during infection **(Supplementary Fig 2A)**. Quantitative investigation of foamy macrophage development provides additional evidence for the qualitative Oil Red O staining results **(Supplementary Figure 2B)**. Macrophages from MDR-PTB patients showed a higher percentage of foamy macrophages under all lipid treatment conditions (naïve LDL, palmitic acid, and oleic acid) in comparison to healthy donors, which is consistent with the increased intracellular lipid accumulation shown in Oil Red O-stained cells. Quantification of lipid accumulated macrophages demonstrates increased foam cell production in MDR-PTB patients. These observations support the Oil Red O staining that shows MDR-PTB patients have higher levels of lipid droplet accumulation and foamy macrophage differentiation, which is related to alterations in lipid metabolism and metabolic reprogramming. The elevated lipid accumulation in monocyte population from MDR patients indicates metabolic reprogramming of monocytes, which may support immunological regulation and bacterial persistence in the host environment. Based on qualitative and quantitative analysis of Oil Red O staining, we predicted changes in nitric oxide levels that correlate with the functional plasticity of macrophages. Since foamy macrophages are believed to be Th2 in nature, these macrophages normally produce less NO compared to normal macrophages, when stimulated in a TLR dependent manner. The observed variations in NO production between the MDR-PTB patients and healthy donors shows difference in immune response state and macrophage activation. Macrophages from TB patients exhibited increased NO levels that indicate impaired antimicrobial activity, which reflects the pathogen’s capacity to alter host defence mechanisms and affects the course of disease **(Supplementary Figure 2C, D)**. Oil red patterning indicates the propensity of lipid loaded macrophages to become foamy macrophages. These macrophages contribute to the pathogenic response in TB patients and exhibit scavenging phenotype.

After the standardization of the protocol, we exposed these foamy macrophages to S1P and analyzed both oil red staining pattern along with key markers Lipid accumulation increased in the healthy donors and MDR-MDR-PTB derived macrophages treated with naïve LDL, oxidized LDL and palmitic acid in which Oil Red O staining and proportion of foamy macrophages increased significantly. Compared to the cells without lipid treatment, the number of lipid droplet was increased in the lipid treated cells. However, the intracellular lipid accumulation and the percentage of foamy macrophages were decreased when both healthy donors and MDR-MDR-PTB derived macrophages treated with S1P **(Supplementary Figure 3A, B, C)**, which means that they had an effect in regulating the lipid homeostasis and attenuated the formation of foam cell **(Figure 2A, B, C)**. Moreover, the production of nitric oxide was elevated in MDR-MDR-PTB patient macrophages compared to healthy donors, however it was not statistically significant **(Figure 3A, B)**. Hence, MDR-MDR-PTB macrophages shown to have higher lipid accumulation and potential tendency to stimulate the inflammatory response, where S1P can efficiently inhibit the burden of lipids.

**Figure 2.**
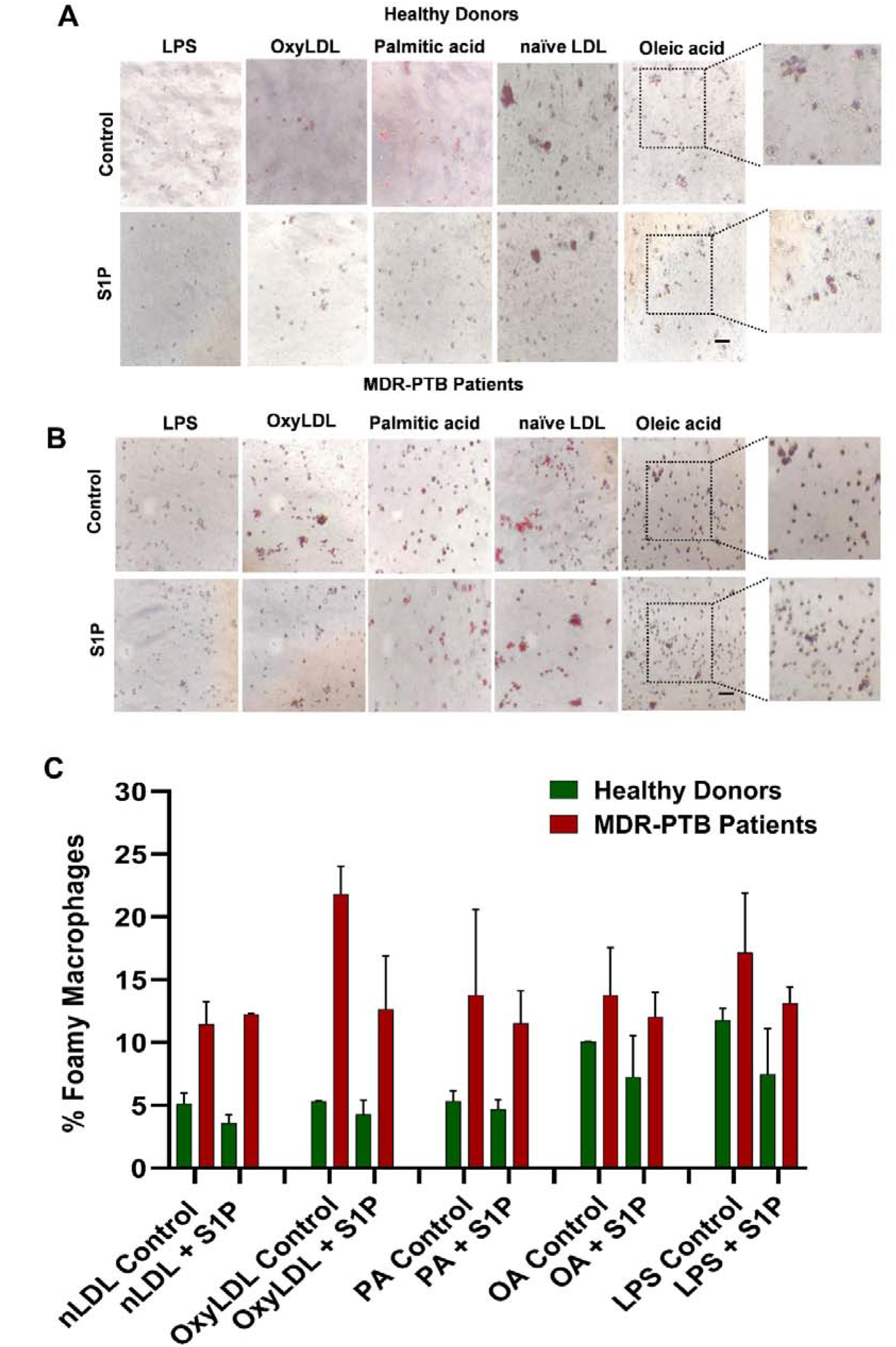
S1P inhibits foam cell formation in primary CD14^+^ monocytes upon lipid treatment. **(A, B)** Representative phase-contrast Oil Red O stained images showing neutral lipid accumulation in **(A)** Healthy donor (n=4) and **(B)** MDR patient (n=4) macrophages treated with LPS, oxyLDL, palmitic acid, naïve LDL, and oleic acid in presence or absence of S1P. **(C)** Quantification of the percent foam macrophages with reduced foam cell formation among different treatment groups especially in OxyLDL and LPS induced in MDR Patients. Data represented as the mean ± SEM.

**Figure 3.**
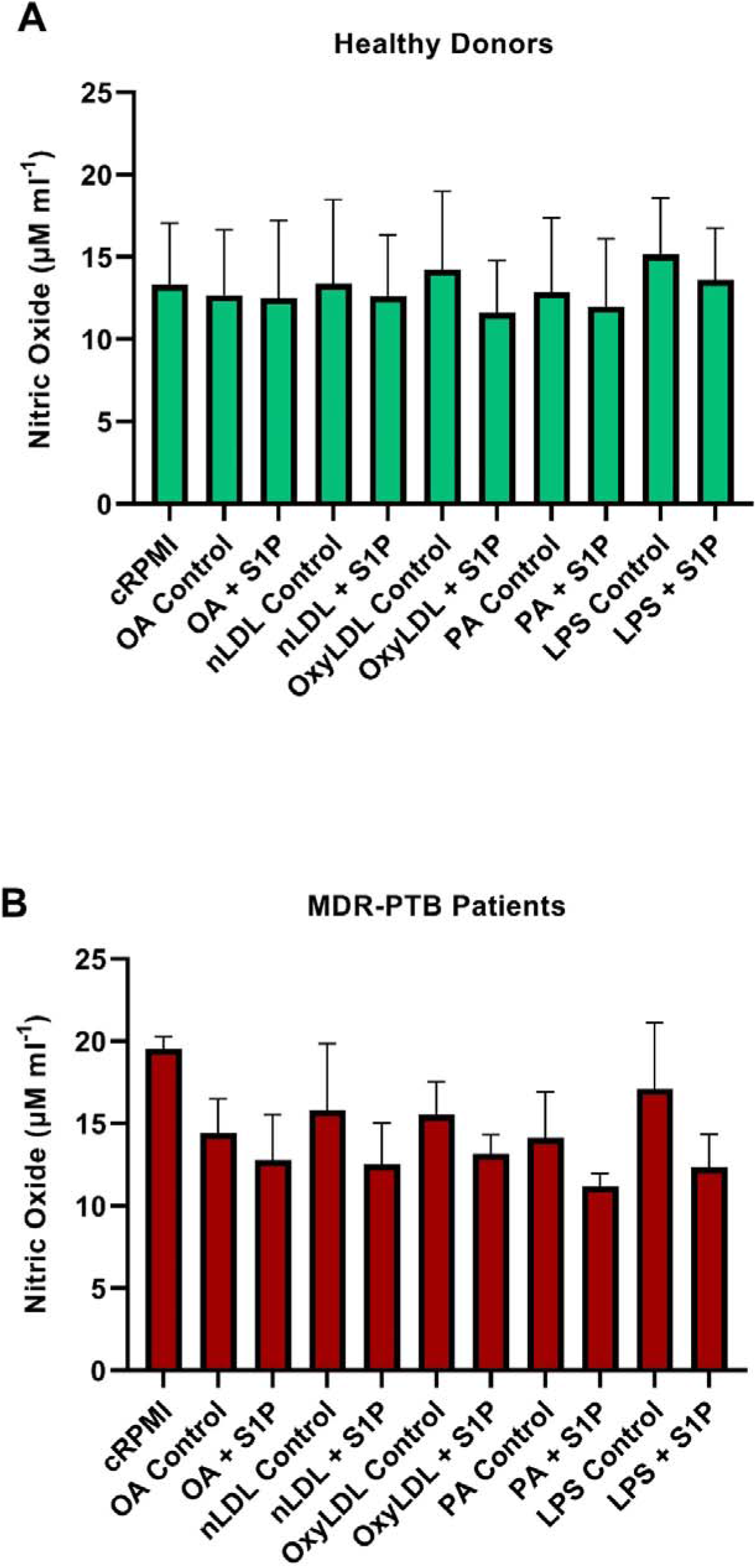
S1P reduces nitric oxide production in CD14^+^ monocytes. **(A, B)** Nitric oxide in culture supernatants of **(A)** Healthy donor (n=4) and **(B)** MDR-PTB patient (n=4) macrophages treated with Oleic acid, naïve LDL, oxyLDL, palmitic acid, and LPS with or without S1P. S1P downregulates NO production in all lipid and inflammatory profiles in MDR-PTB macrophages whereas Healthy donor macrophages show a more stable NO production. Data represented as the mean ± SEM.

### Lipid Metabolic and ER stress genes exhibit Differential Expression in MDR-MDR-PTB Macrophages

Following lipid treatments, RT-PCR analysis showed differential expression of genes linked to lipid metabolism, immune modulation, and ER stress. Both healthy donor and MDR-PTB macrophages showed elevated *APOE* expression, especially following oleic acid treatment. In response to lipid stimuli, *LDLR1* exhibited modest upregulation, indicating improved lipid absorption. The level of *CLEC7A* was significantly high in MDR-PTB macrophages, while *CLEC4E* exhibited differential expression across the treatment groups. With respect to the immune-regulatory genes, the expressions of *IL-4* and *IL-13* were slightly upregulated in MDR-PTB macrophages, as compared to healthy individuals, suggesting that an anti-inflammatory state is favoured. The expression level of *GP96* was moderately increased in MDR-PTB macrophages, suggesting the induction of ER stress response **(Figure 4, 5)**. The obtained data revealed that lipid treatment could induce transcriptomic changes, corresponding to lipid accumulation, macrophage polarization and immune regulation, especially for MDR-MDR-PTB individuals.

**Figure 4.**
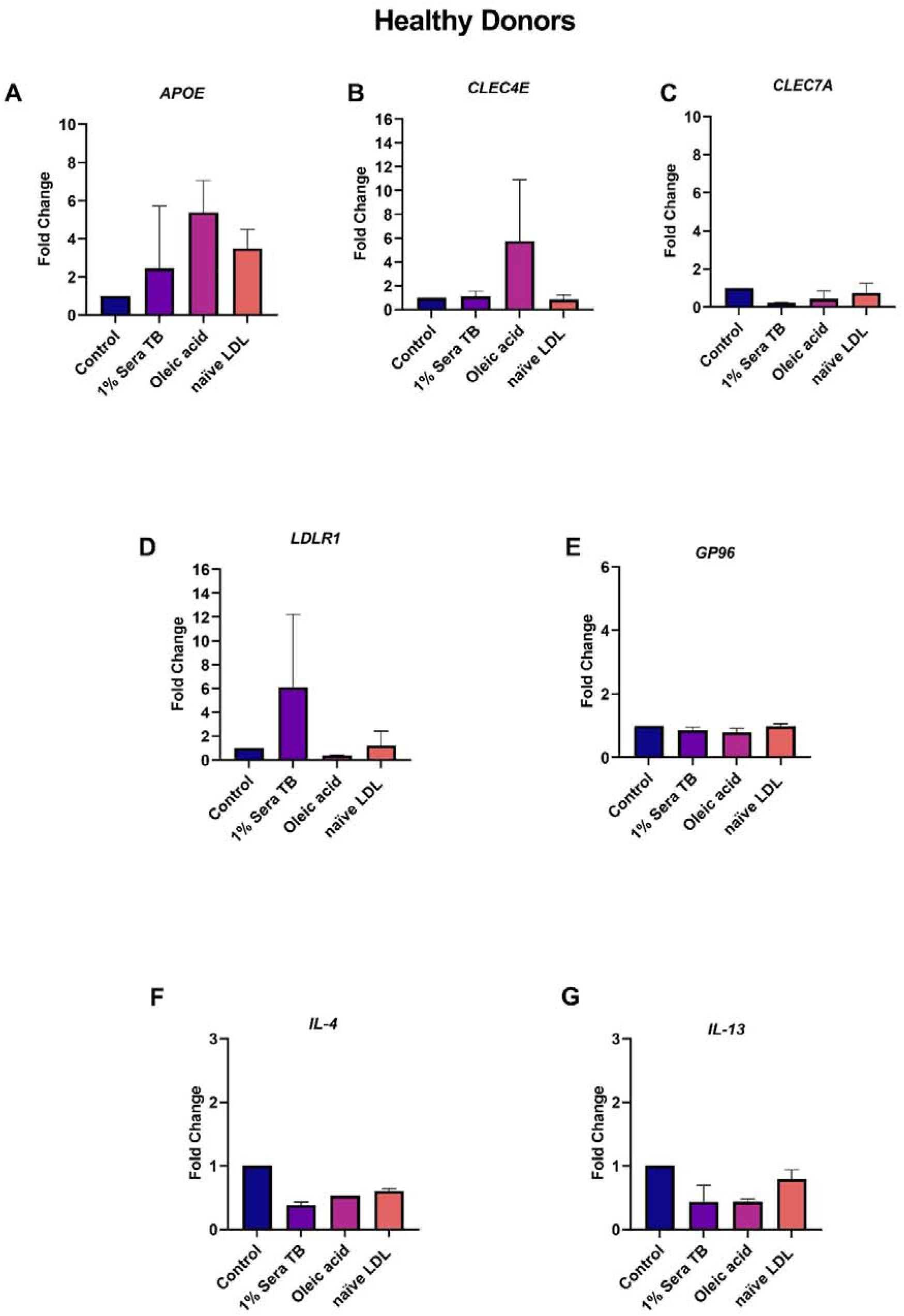
Oleic acid and TB serum differentially affect lipid metabolism and C-type lectin receptor gene expression in healthy donor macrophages. **(A–G)** mRNA expression fold change of (A) *APOE*, (B) *CLEC4E* (Mincle), (C) *CLEC7A* (Dectin-1), (D) *LDLR1*, (E) *GP96*, (F) *IL-4* and (G) *IL-13* of Healthy donor macrophages stimulated with Control, 1% Sera TB, Oleic acid and naïve LDL. Oleic acid selectively upregulates *APOE* and *CLEC4E* while 1% Sera TB strongly upregulates *LDLR1* expression. Fold change was calculated using the 2^-ΔΔ^Ct method and normalized to β-actin. Data represent gene expression profiles obtained from two healthy donors.

**Figure 5.**
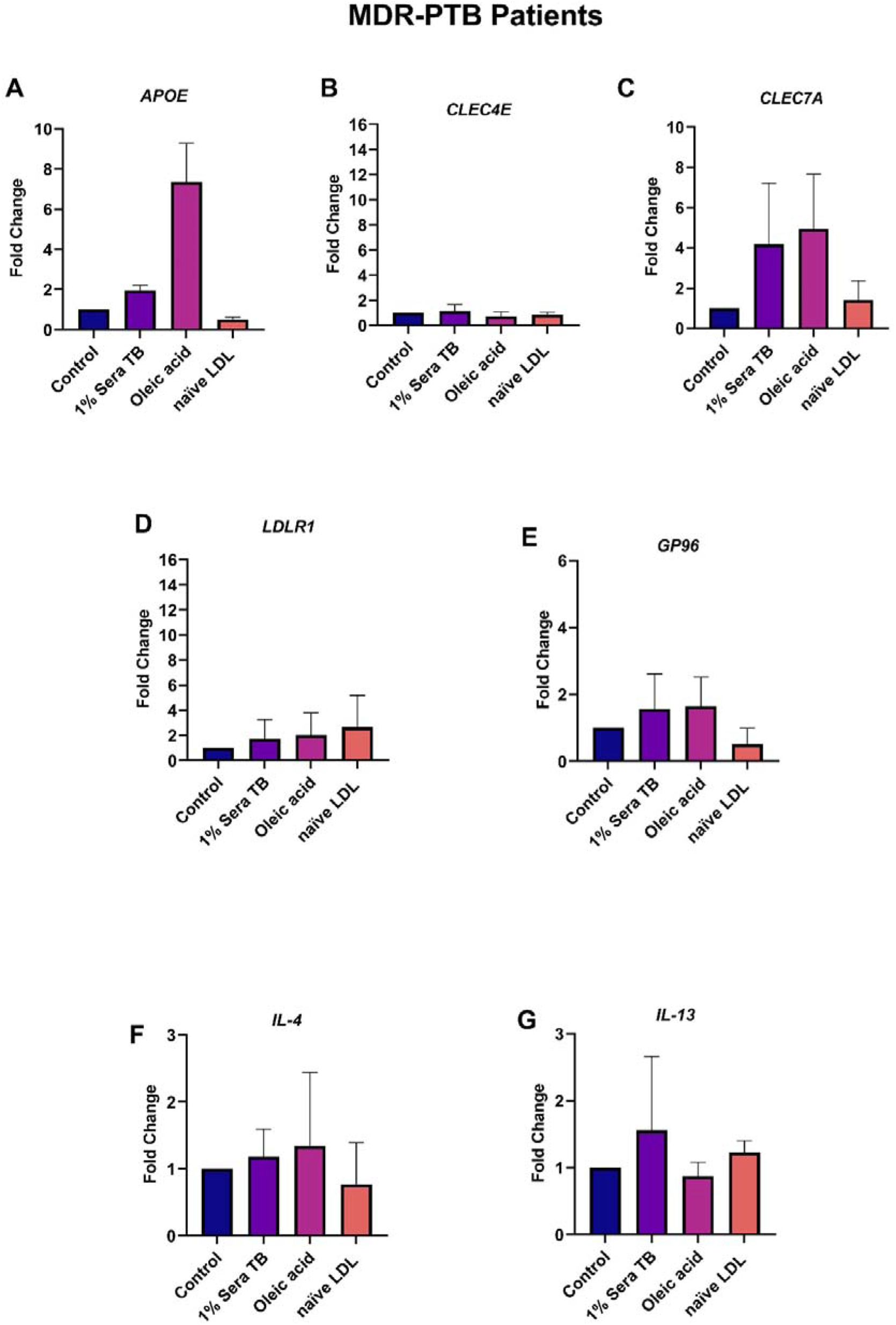
Oleic acid and TB serum selectively upregulates *APOE* and *CLEC7A* in MDR-PTB patient macrophages. **(A–G)** The mRNA expressions of (A) *APOE*, (B) *CLEC4E*, (C) *CLEC7A* (Dectin-1), (D) *LDLR1*, (E) *GP96*, (F) *IL-4* and (G) *IL-13* were determined as fold change in macrophages from MDR-PTB patients treated with Control, 1% Sera TB, Oleic acid or naïve LDL. Oleic acid upregulates the gene expression of *APOE* (∼7-fold), *CLEC7A* (∼5-fold), and 1% Sera TB has an upregulation restricted to *CLEC7A* (∼4-fold) with reduced levels of induction for *CLEC4E* compared with Healthy Donors. Fold change was calculated using the 2-ΔΔCt method and normalized to β-actin. Data represent gene expression profiles obtained from two MDR patients.

S1P treated macrophages derived from healthy donors resulted in significantly decreased *APOE* expression following oleic acid treatment, whereas in the naïve LDL treated group increased expression was observed. *LDLR1* and *GP96* showed an increasing trend following oleic acid treatment whereas *IL-4* expression increased slightly and *IL-13* decreased **(Figure 6)**. S1P treated macrophages derived from MDR-MDR-PTB patients exhibited increased *CLEC4E* and *IL-13* expression following oleic acid and naïve LDL treatment respectively. *APOE*, *LDLR1*, *GP96* and *IL-4* expression varied across treatment groups **(Figure 7)**. These results suggest that S1P treatment alters the expression of genes related to lipid and immune regulation variation in macrophages from healthy donors and in MDR-MDR-PTB patients.

**Figure 6.**
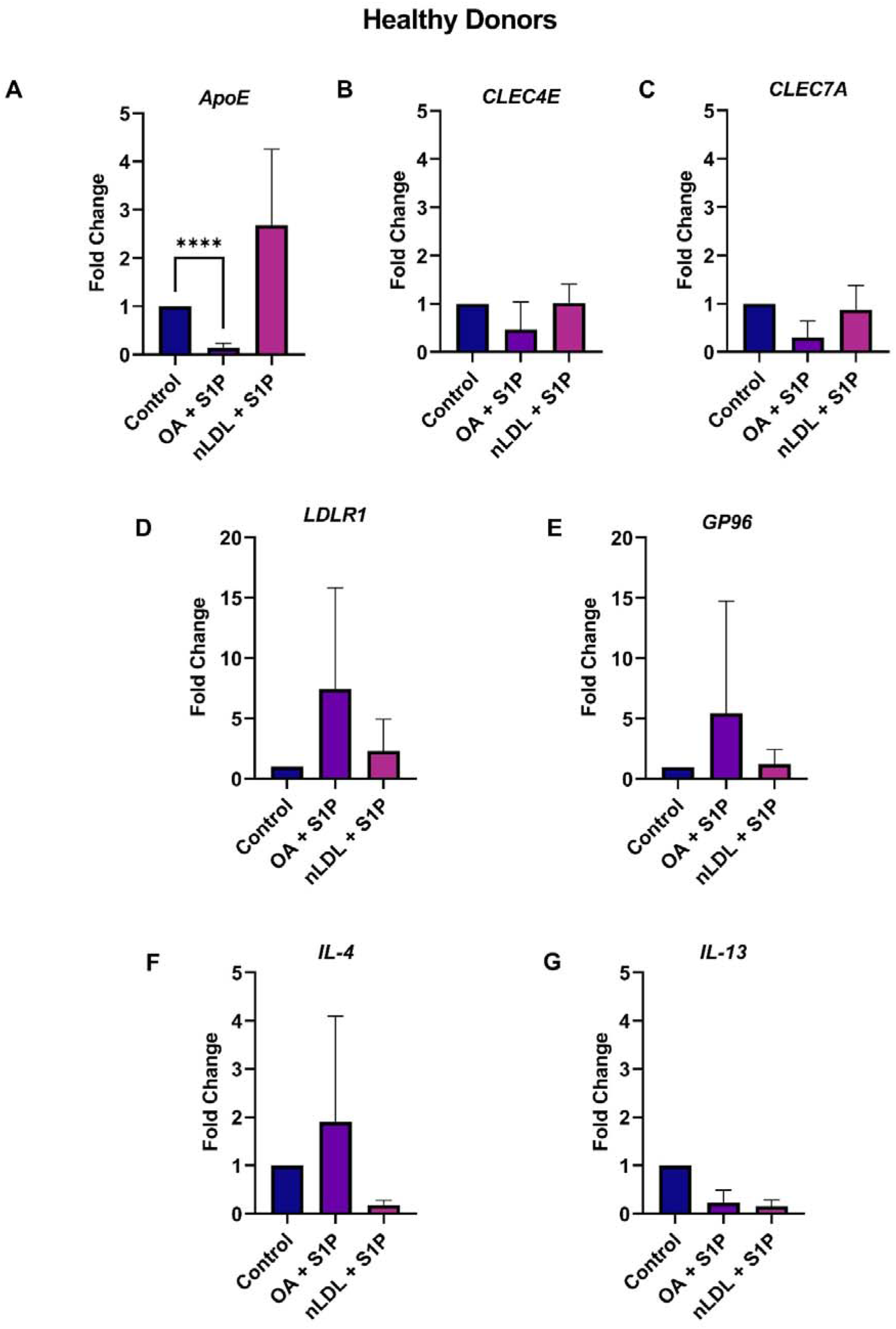
Gene expression after S1P treatment in macrophages from healthy donors. **(A-G)** Fold change in relative gene expression of *APOE*, *CLEC4E*, *CLEC7A*, *LDLR1*, *GP96*, *IL-4* and *IL-13* following exposure to oleic acid and naïve LDL, then subsequent S1P treatment, in comparison with the control. Data show fold change over control. One-way Anova test has been performed. Statistical significance (****p<0.0001), (n=4).

**Figure 7.**
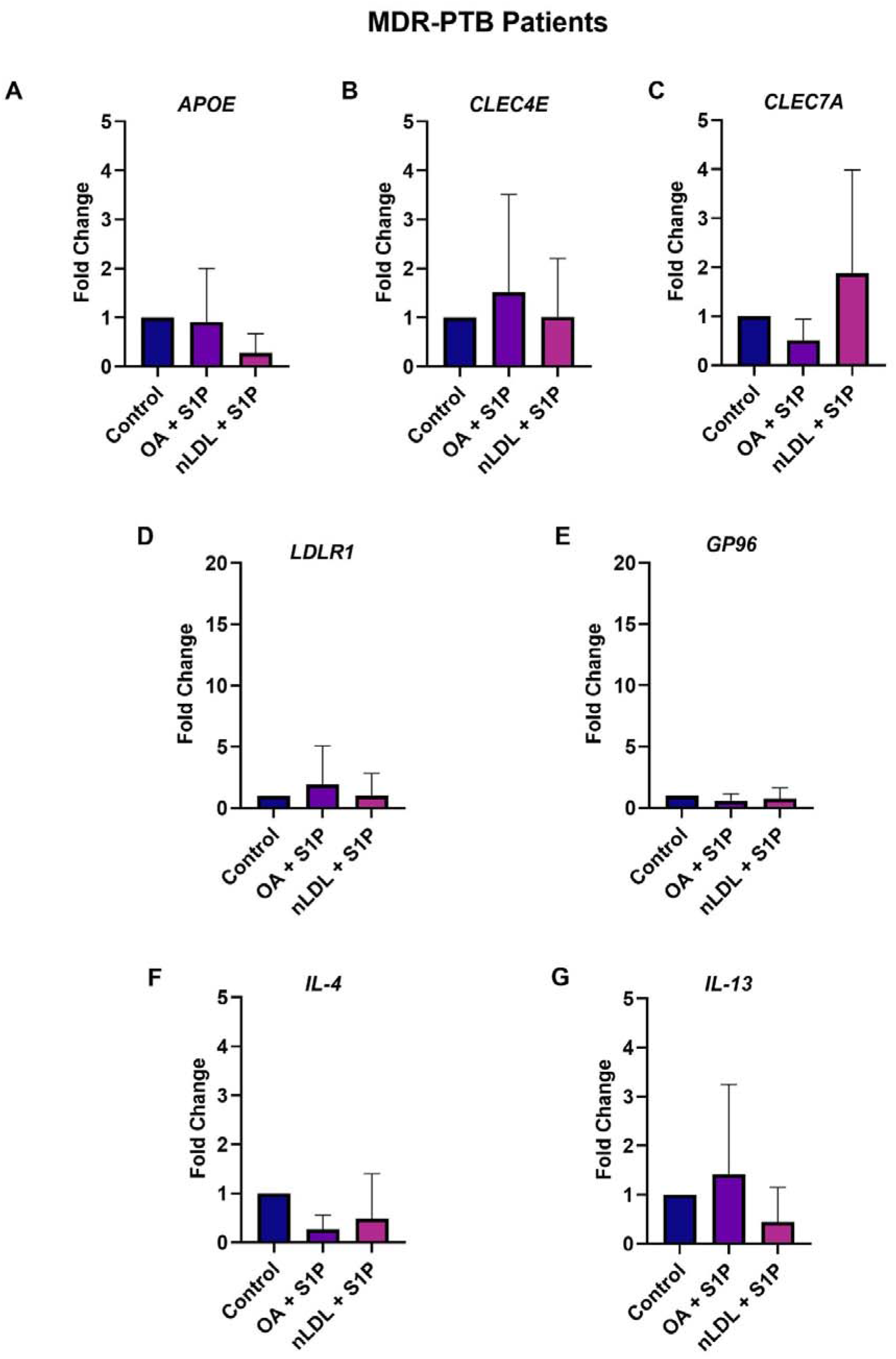
Gene expression in MDR-PTB patient-derived macrophages following S1P treatment. **(A-G)** Expression of *APOE*, *CLEC4E*, *CLEC7A*, *LDLR1*, *GP96*, *IL-4* and *IL-13* in response to oleic acid and naïve LDL exposure and subsequent treatment with S1P. Results are shown as the fold change in expression relative to the control (n=4).

## Discussion

S1P and low-density lipoprotein (LDL) work together and influence each other, playing a central role in signalling during pathogenic inflammatory response. Polyunsaturated fatty acids (PUFAs) and other types of structural and dietary fats also help to regulate this process. An interesting fact regarding the S1P receptor biology is the difference in receptor affinity between species: although human naïve signalling occurs mainly through the S1PR1 receptor, whereas S1P binds to the S1PR3 receptor in mouse models (5, 30). At the cellular level, S1P signalling regulates the PI3K pathway, and calcium (Ca2+) transport from the ER, clarifying how this specific activity undermines LDL homeostasis and lipid regulation (31). Specifically related to the immunomodulation, there is a strong association between S1P signalling and Th1 responses, demonstrating the crosstalk among these lipids (5). Among membrane receptors, CLEC7A (Dectin-1) helps bring together these crosstalk mechanisms to manage cellular lipid balance (32). The production of “foamy” macrophages (FM) is a key biological characteristic of TB. The term “foamy” was first used to describe the frothy appearance of lipid-filled macrophages in atherosclerotic disease. Chloroquine, a medication that prevents acidification to inactivate lysosomal hydrolases, such as lysosomal acid lipase (LAL), which is necessary for the catalysis of cholesterol esters, may be used in experiments to produce foamy cells (33). Glycerophospholipids (GPs), glycerolipids, sphingolipids, sterols, fatty acids (FAs), prenols, saccharolipids, and polyketides are the eight types of lipids that together comprise cellular membranes (34). Cholesterol is the major source of carbon during the infection process of M.tb. Cholesterol is important for the proper growth and persistence of *M.tb*. It acts as an essential growth substrate for *M.tb* inside phagosomes and facilitates the formation of foamy macrophages and necrotic granulomas. *M.tb* can utilize cholesterol as its only carbon source for energy production and membrane lipid formation, which in turn leads to the synthesis of virulence lipids in its cell wall (35). Although mycolic acids do not appear to make up the majority of the lipids in foam cells, they may contribute to the growth and functioning of foam cells by controlling genes linked to lipids that are downstream of Toll-like receptors (TLRs) and scavenger receptor pathways. For instance, human and murine macrophage cytokine responses to M.tb and mycobacterial trehalose dimycolate require TLR2, and CD14 (36). Foamy macrophages can develop under different pathological conditions. They are not limited to atherosclerosis; foam cells are a characteristic of tuberculosis infection and caseation, which is a sign of ongoing mycobacterial immune responses. Mycobacterial granulomata include a large number of foam cells. It is generally believed that these cells accumulate lipids and eventually die, causing caseation, which exacerbates tissue damage, chronic inflammation, and the pathogen’s dissemination (20). Foamy macrophage production is facilitated by numerous ways that *M.tb* and bacterial factors alter macrophage lipid metabolism. For instance, ESAT-6 increases lipid synthesis and improves glucose metabolism in macrophages by activating the G-protein-coupled receptor (37).

Our study nicely demonstrates that macrophages derived from PBMCs of MDR-PTB patients showed strong metabolic reprogramming as compared to healthy control macrophages. While both groups of PBMCs differentiated well into mature macrophages when cultured in M-CSF, PBMCs from MDR-PTB patients exhibited more prominent foamy morphology and intracellular lipid accumulation. Consequently, these findings indicate that in MDR-PTB, there is a remodelling of lipid metabolism that results in a lipid-rich intracellular environment, potentially facilitating bacterial persistence. Lipid accumulation in macrophages is relatively well described in the context of atherosclerosis but is poorly understood in the context of *M.tb* infection, especially in the context of MDR-PTB infection. Thus, native LDL, oxidized LDL and palmitic acid induced immense accumulation of lipids and foam cell transformation of macrophages, whereas S1P treatment was able to reduce the intracellular lipid content in macrophages of both patient groups, being more effective in MDR-PTB patients.

These results suggest that S1P alters lipid homeostasis in macrophages, potentially countering the metabolic reprogramming that characterizes foamy macrophages. There was higher NO production in macrophages infected with MDR-PTB compared to macrophages from healthy donors, although this was not significant possibly due to the high variability in clinical data. These findings suggest that MDR-PTB macrophages potentially decouple immune-metabolic arm response. Consistent with these findings, we observe dysregulation of lipid metabolism as well as immune-regulatory genes (e.g. *APOE, LDLR1, CLEC4E, CLEC7A, GP96, IL-4* and *IL-13*), likely reflecting a strong interconnection in metabolic and immune reprogramming, which can also influence the polarization of macrophages. This study is the first study which highlights the significance of host sphingolipids in disrupting the integrity of foamy macrophages and opens new window for a host directed approach against MDR-PTB.

## Acknowledgement

We thank all the study participants and the staff members from Department of Pulmonary Medicine, All India Institute of Medical Sciences, New Delhi. We acknowledge Dr. Nilanjana Basu, Amity Institute of Molecular Medicine & Stem Cell Research, and Dr. Amrita Pathak, Amity Institute of Neuropsychology & Neurosciences, Amity University for providing help in real time experiments.

## Data Availability

The data supporting the findings of this study are available from the corresponding author upon reasonable request. Data generated from PBMC derived Macrophages isolated from healthy donors and patients with MDR-PTB/EMDR-PTB are not publicly available because they contain sensitive human participant information.

## Author Contributions

H.P. and V.H. conceived and designed the study and planned the experiments. A.K. and N.S. performed the experiments. V.H., L.K.S. and A.K.S. supervised patient recruitment and the prospective clinical study involving CD14 monocytes from healthy donors and MDR-PTB patients and contributed to manuscript editing. H.P., A.K.S., A.K., V.H. and L.K.S. conducted data analysis, visualization of data, and provided methodological guidance. H.P., A.K. and L.K.S. prepared the original draft of the manuscript. All authors reviewed, revised, and approved the final version of the manuscript.

## Supplementary File

### Supplementary Figures

**Suppl Figure 1.**
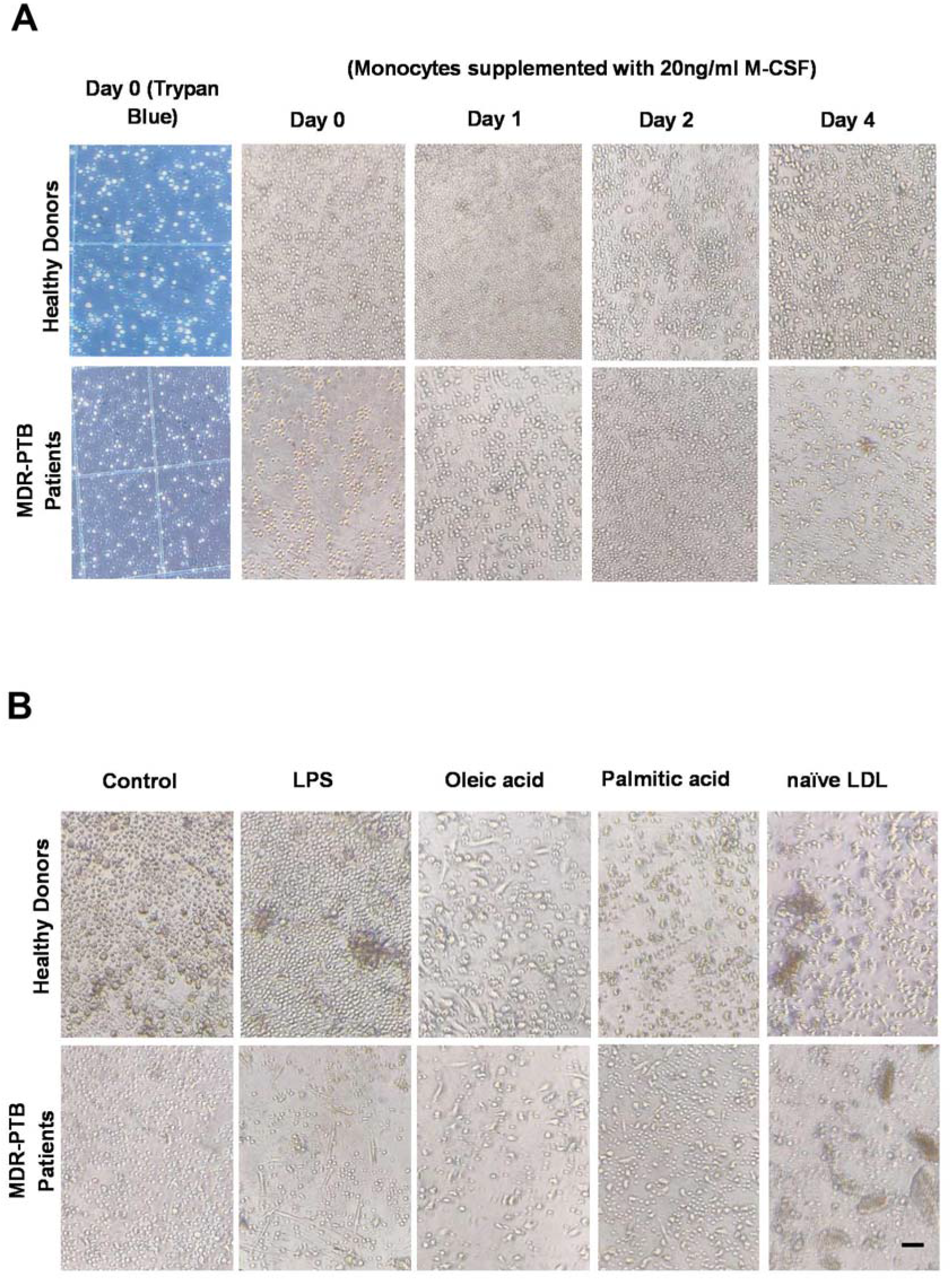
Macrophage Differentiation and Morphology of the Macrophage upon Lipid Treatment. **(A)** Differentiation of primary human monocytes (Healthy Donors and MDR Patients) with 20ng/ml M-CSF at Day 0-4 (time-course differentiation). **(B)** Phase-contrast images of Day 4 macrophages (Control, LPS, OA, PA and naive LDL treatment).

**Suppl Figure 2.**
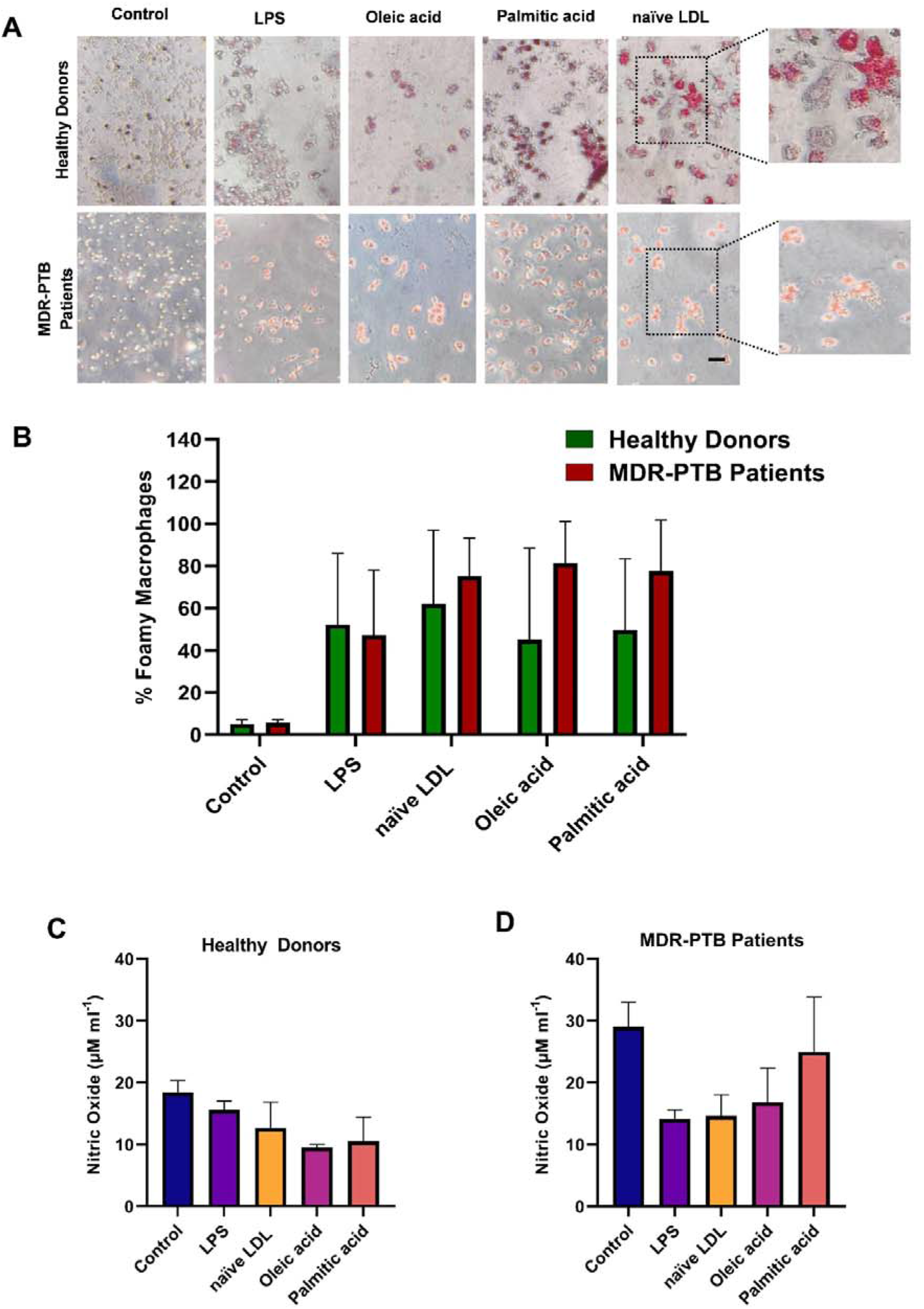
Differential foam cell formation among MDR patients and healthy donors CD14^+^ monocytes. **(A)** Representative images of macrophages stained with Oil Red O showing neutral lipid accumulation following treatment with control, LPS, oleic acid, palmitic acid and naïve LDL in Healthy donors (HD, n=3) and Multidrug-Resistant Tuberculosis (MDR-TB, n=3) patients. **(B)** Percentage of foamy macrophages was quantified in each treatment groups. Data represented as the mean ± SEM. **(C, D)** Nitric oxide measured in culture supernatants from **(A)** Healthy donor (n=3) and **(B)** MDR-TB patient (n=3) macrophages following different lipid treatments. Data represented as the mean ± SEM.

**Suppl Fig. 3.**
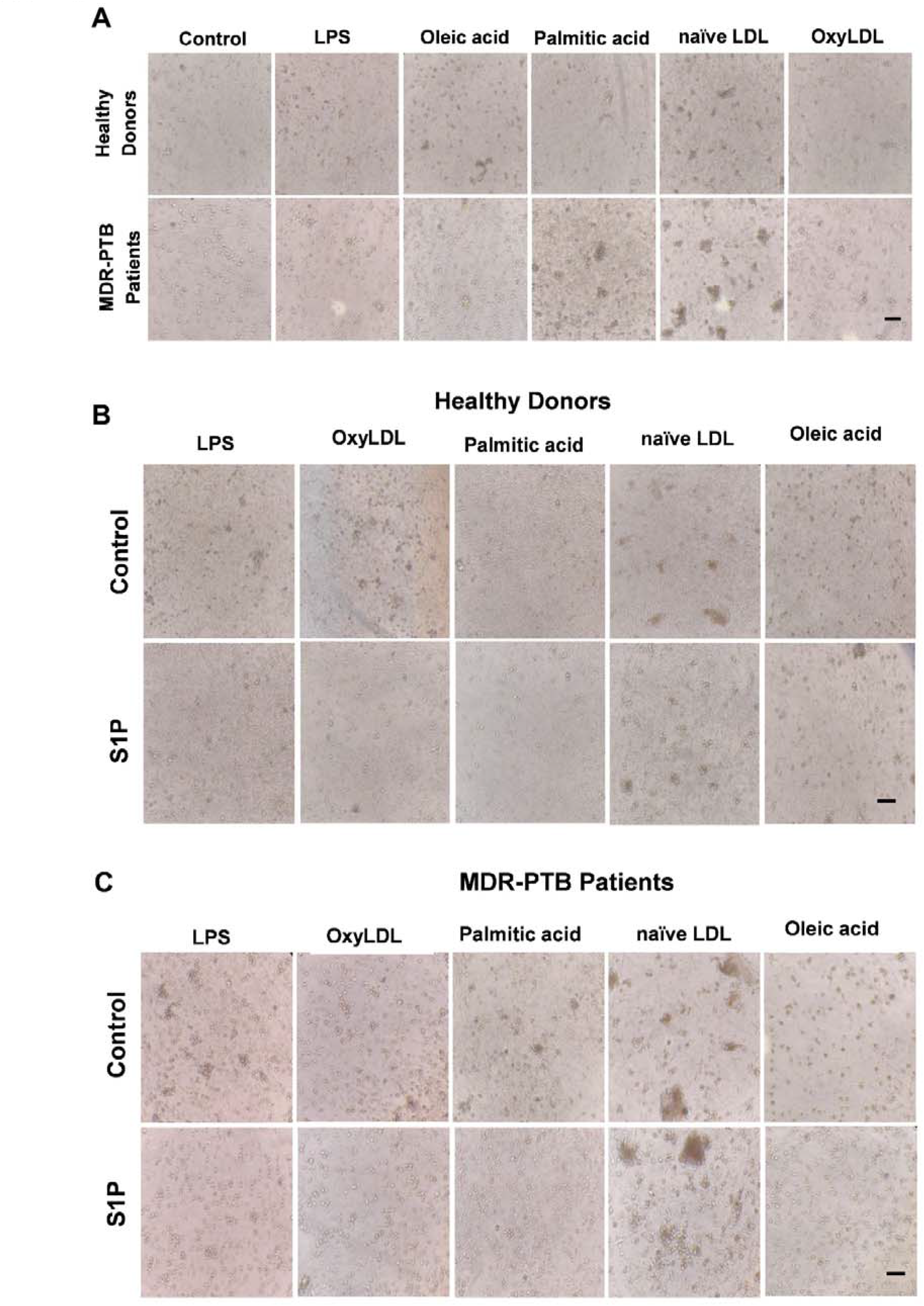
S1P regulates Morphological Changes in Macrophages Induced by Lipids and LPS. **(A)** Phase-contrast images of Day 4 Healthy Donor and MDR Patient macrophages treated with Control, LPS, Oleic acid, Palmitic acid, naive LDL and oxyLDL. **(B, C)** Morphological analysis of **(B)** Healthy Donor and **(C)** MDR Patient macrophages treated with different lipids or LPS with or without S1P.

**Supplementary Table 1.**
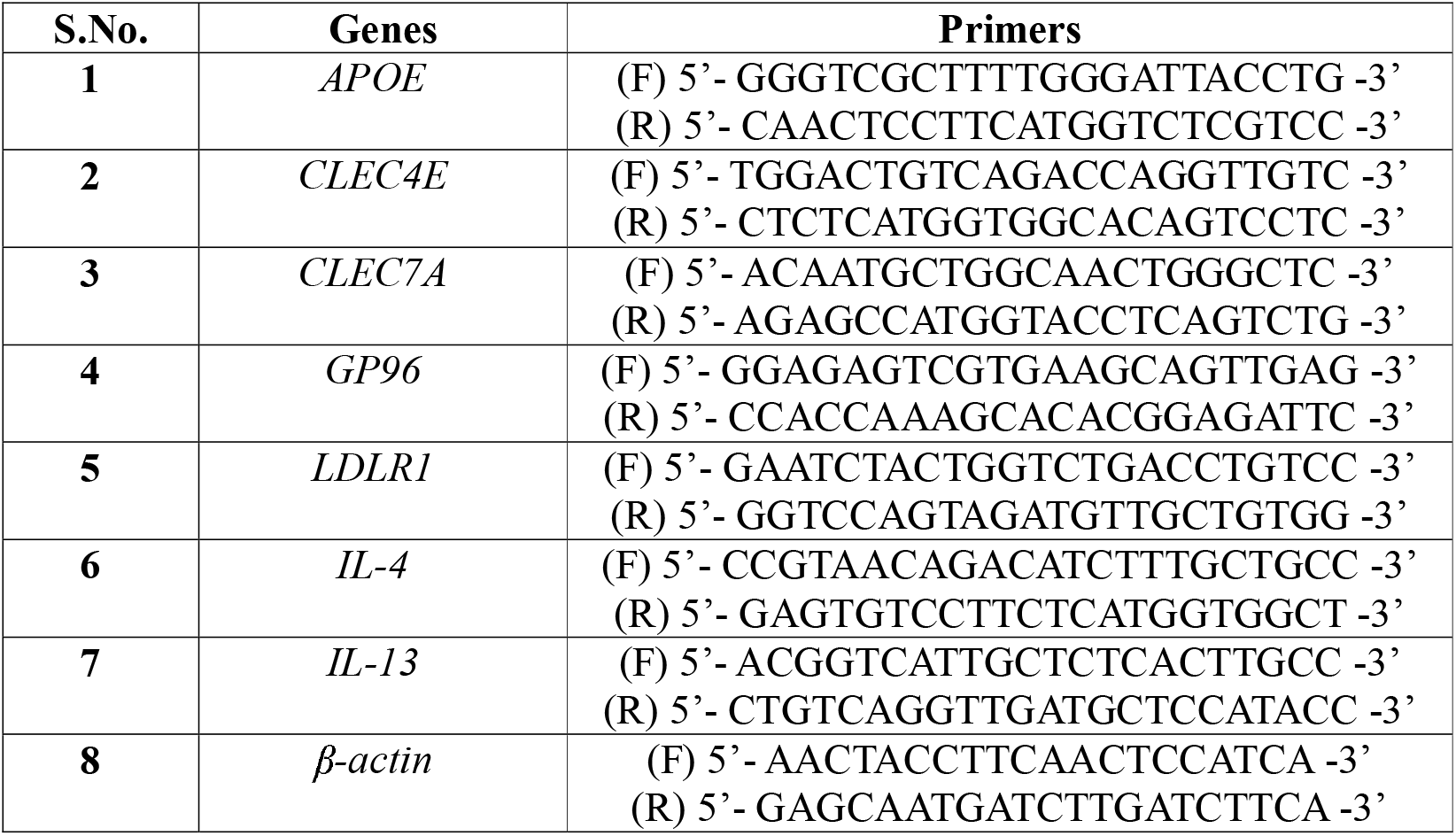
Primers used in this study.

## References

1. Lanni, F., Wijnant, G.J., Xie, M., Osiecki, P., Dartois, V. and Sarathy, J.P., 2023. Adaptation to the intracellular environment of primary human macrophages influences drug susceptibility of *Mycobacterium tuberculosis*. Tuberculosis, 139, p.102318. doi: 10.1016/j.tube.2023.102318.

2. Kirubakar, G., Johnston, L., Nae Rin Lee, B., Russell, D.G. and Simwela, N.V., 2025. Incorporation of macrophage immune stresses into an intracellular assay of drug tolerance in *Mycobacterium tuberculosis*. Antimicrobial Agents and Chemotherapy, 69(10), p.e00795–25. doi: 10.1128/aac.00795-25.

3. Ufimtseva, E.G. and Eremeeva, N.I., 2023. Drug-tolerant *Mycobacterium tuberculosis* adopt different survival strategies in alveolar macrophages of patients with pulmonary tuberculosis. International Journal of Molecular Sciences, 24(19), p.14942. doi: 10.3390/ijms241914942.

4. Santucci, P., Greenwood, D.J., Fearns, A., Chen, K., Jiang, H. and Gutierrez, M.G., 2021. Intracellular localisation of *Mycobacterium tuberculosis* affects efficacy of the antibiotic pyrazinamide. Nature communications, 12(1), p.3816. doi: 10.1038/s41467-021-24127-3.

5. Russell, D.G., Cardona, P.J., Kim, M.J., Allain, S. and Altare, F., 2009. Foamy macrophages and the progression of the human tuberculosis granuloma. Nature Immunology, 10(9), p.943–948. doi: 10.1038/ni.1781.

6. Kim, M.J., Wainwright, H.C., Locketz, M., Bekker, L.G., Walther, G.B., Dittrich, C., Visser, A., Wang, W., Hsu, F.F., Wiehart, U. and Tsenova, L., 2010. Caseation of human tuberculosis granulomas correlates with elevated host lipid metabolism. EMBO Molecular Medicine, 2(7), p.258–274. doi: 10.1002/emmm.201000079.

7. Shim, D., Kim, H. and Shin, S.J., 2020. *Mycobacterium tuberculosis* infection-driven foamy macrophages and their implications in tuberculosis control as targets for host-directed therapy. Frontiers in Immunology, 11, p.910. doi: 10.3389/fimmu.2020.00910.

8. Laval, T., Chaumont, L. and Demangel, C., 2021. Not too fat to fight: The emerging role of macrophage fatty acid metabolism in immunity to *Mycobacterium tuberculosis*. Immunological Reviews, 301(1), p.84–97. doi: 10.1111/imr.12952.

9. Peyron, P., Vaubourgeix, J., Poquet, Y., Levillain, F., Botanch, C., Bardou, F., Daffé, M., Emile, J.F., Marchou, B., Cardona, P.J. and De Chastellier, C., 2008. Foamy macrophages from tuberculous patients’ granulomas constitute a nutrient-rich reservoir for *M. tuberculosis* persistence. PLoS Pathogens, 4(11), p.e1000204. doi: 10.1371/journal.ppat.1000204.

10. Marino, S., Cilfone, N.A., Mattila, J.T., Linderman, J.J., Flynn, J.L. and Kirschner, D.E., 2015. Macrophage polarization drives granuloma outcome during *Mycobacterium tuberculosis* infection. Infection and Immunity, 83(1), p.324–338. doi: 10.1128/IAI.02494-14.

11. Liu, Y., Tan, S., Huang, L., Abramovitch, R.B., Rohde, K.H., Zimmerman, M.D., Chen, C., Dartois, V., VanderVen, B.C. and Russell, D.G., 2016. Immune activation of the host cell induces drug tolerance in *Mycobacterium tuberculosis* both in vitro and in vivo. Journal of Experimental Medicine, 213(5), p.809–825. doi: 10.1084/jem.20151248.

12. Wallis, R.S. and Hafner, R., 2015. Advancing host-directed therapy for tuberculosis. Nature Reviews Immunology, 15(4), p.255–263. doi: 10.1038/nri3813.

13. Nadella, V., Sharma, L., Kumar, P., Gupta, P., Gupta, U.D., Tripathi, S., Pothani, S., Qadri, S.S.Y.H. and Prakash, H., 2020. Sphingosine-1-phosphate (S-1P) promotes differentiation of naïve macrophages and enhances protective immunity against *Mycobacterium tuberculosis*. Frontiers in Immunology, 10, p.3085. doi: 10.3389/fimmu.2019.03085.

14. S.K., Volpe, E., Palmieri, G., Mattei, M., Galati, D., Martino, A., Piccioni, M.S., Valente, E., Bonanno, E., De Vito, P. and Baldini, P.M., 2004. Sphingosine 1–phosphate induces antimicrobial activity both in vitro and in vivo. The Journal of Infectious Diseases, 189(11), p.2129–2138. doi: 10.1086/386286.

15. Yadav, V., Sahoo, S., Malhotra, N., Mishra, R., Sreedharan, S., Rajmani, R.S., Shanmugam, S., Shandil, R.K., Narayanan, S., Thacker, V.V. and Laxman, S., 2025. Bioenergetic reprogramming of macrophages reduces drug tolerance in *Mycobacterium tuberculosis*. Nature Communications, 16(1), p.9370. doi: 10.1038/s41467-025-64407-w.

16. Arish, M. and Naz, F., 2022. Sphingosine-1-phosphate receptors 2 and 3 reprogram resting human macrophages into M1 phenotype following mycobacteria infection. Current Research in Immunology, 3, p.110–117. doi: 10.1016/j.crimmu.2022.05.004.

17. Weigert, A., Weis, N. and Brüne, B., 2009. Regulation of macrophage function by sphingosine-1-phosphate. Immunobiology, 214(9-10), p.748–760. doi: 10.1016/j.imbio.2009.06.003.

18. Bowman, E.R., Cameron, C.M., Richardson, B., Kulkarni, M., Gabriel, J., Cichon, M.J., Riedl, K.M., Mustafa, Y., Cartwright, M., Snyder, B. and Raman, S.V., 2020. Macrophage maturation from blood monocytes is altered in people with HIV, and is linked to serum lipid profiles and activation indices: A model for studying atherogenic mechanisms. PLoS Pathogens, 16(10), p.e1008869. doi: 10.1371/journal.ppat.1008869.

19. Hamidzadeh, K., Belew, A.T., El-Sayed, N.M. and Mosser, D.M., 2020. The transition of M-CSF–derived human macrophages to a growth-promoting phenotype. Blood Advances, 4(21), p.5460–5472. doi: 10.1182/bloodadvances.2020002683.

20. Agarwal, P., Combes, T.W., Shojaee-Moradie, F., Fielding, B., Gordon, S., Mizrahi, V. and Martinez, F.O., 2020. Foam cells control *Mycobacterium tuberculosis* infection. Frontiers in Microbiology, 11, p.1394. doi: 10.3389/fmicb.2020.594142.

21. Caocci, M., Niu, M., Fox, H.S. and Burdo, T.H., 2024. HIV infection drives foam cell formation via NLRP3 inflammasome activation. International Journal of Molecular Sciences, 25(4), p.2367. doi: 10.3390/ijms25042367.

22. Bruggen, van der T., Nijenhuis, S., van Raaij, E., Verhoef, J. and Sweder van Asbeck, B., 1999. Lipopolysaccharide-induced tumor necrosis factor alpha production by human monocytes involves the raf-1/MEK1-MEK2/ERK1-ERK2 pathway. Infection and Immunity, 67(8), p.3824–3829 doi: 10.1128/IAI.67.8.3824-3829.1999.

23. Al-Rashed, F., Haddad, D., Al Madhoun, A., Sindhu, S., Jacob, T., Kochumon, S., Obeid, L.M., Al-Mulla, F., Hannun, Y.A. and Ahmad, R., 2023. ACSL1 is a key regulator of inflammatory and macrophage foaming induced by short-term palmitate exposure or acute high-fat feeding. iScience, 26(7). doi: 10.1016/j.isci.2023.107145.

24. Ramachandran, S., Vinitha, A. and Kartha, C.C., 2016. Cyclophilin A enhances macrophage differentiation and lipid uptake in high glucose conditions: a cellular mechanism for accelerated macro vascular disease in diabetes mellitus. Cardiovascular diabetology, 15(1), p.152. doi: 10.1186/s12933-016-0467-5.

25. Sanda, G.M., Stancu, C.S., Deleanu, M., Toma, L., Niculescu, L.S. and Sima, A.V., 2021. Aggregated LDL turn human macrophages into foam cells and induce mitochondrial dysfunction without triggering oxidative or endoplasmic reticulum stress. PLoS One, 16(1), p.e0245797. doi: 10.1371/journal.pone.0245797.

26. Du, J., Zhao, L., Kang, Q., He, Y. and Bi, Y., 2023. An optimized method for Oil Red O staining with the salicylic acid ethanol solution. Adipocyte, 12(1), p.2179334. doi: 10.1080/21623945.2023.2179334.

27. Gap-Gaupool, B., Glenn, S.M., Milburn, E., Turapov, O., Crosatti, M., Hincks, J., Stewart, B., Bacon, J., Kendall, S.L., Voskuil, M.I. and Riabova, O., 2024. Nitric oxide induces the distinct invisibility phenotype of *Mycobacterium tuberculosis*. Communications Biology, 7(1), p.1206. doi: 10.1038/s42003-024-06912-0.

28. Bhakta, S.A., Borba, R., Taba Jr, M., Garcia, C.D. and Carrilho, E., 2014. Determination of nitrite in saliva using microfluidic paper-based analytical devices. Analytica Chimica acta, 809, p.117–122. doi: 10.1016/j.aca.2013.11.044.

29. Luque, M.C.A., Gutierrez, P.S., Debbas, V., Martins, W.K., Puech-Leao, P., Porto, G., Coelho, V., Boumsell, L., Kalil, J. and Stolf, B., 2013. Phage display identification of CD100 in human atherosclerotic plaque macrophages and foam cells. PLoS One, 8(9), p.e75772. doi: 10.1371/journal.pone.0075772. doi: 10.1371/journal.pone.0075772.

30. Borodzicz-Jażdżyk, S., Jażdżyk, P., Łysik, W., Cudnoch-J drzejewska, A. and Czarzasta, K., 2022. Sphingolipid metabolism and signaling in cardiovascular diseases. Frontiers in Cardiovascular Medicine, 9, p.915961. doi: 10.3389/fcvm.2022.915961.

31. Sukhorukov, V.N., Khotina, V.A., Bagheri Ekta, M., Ivanova, E.A., Sobenin, I.A. and Orekhov, A.N., 2020. Endoplasmic reticulum stress in macrophages: the vicious circle of lipid accumulation and pro-inflammatory response. Biomedicines, 8(7), p.210. doi: 10.3390/biomedicines8070210.

32. Tolksdorf, C., Moritz, E., Wolf, R., Meyer, U., Marx, S., Bien-Möller, S., Garscha, U., Jedlitschky, G. and Rauch, B.H., 2022. Platelet-derived S1P and its relevance for the communication with immune cells in multiple human diseases. International Journal of Molecular Sciences, 23(18), p.10278. doi: 10.3390/ijms231810278.

33. Bedard, M., van der Niet, S., Bernard, E.M., Babunovic, G., Cheng, T.Y., Aylan, B., Grootemaat, A.E., Raman, S., Botella, L., Ishikawa, E. and O’Sullivan, M.P., 2023. A terpene nucleoside from *M. tuberculosis* induces lysosomal lipid storage in foamy macrophages. The Journal of Clinical Investigation, 133(6). doi: 10.1172/JCI161944.

34. Seo, J.W., Park, K.S., Lee, G.B., Park, S.E., Choi, J.H. and Moon, M.H., 2023. Comprehensive lipid profiling recapitulates enhanced lipolysis and fatty acid metabolism in intimal foamy macrophages from murine atherosclerotic aorta. Immune network, 23(4), p.e28. doi: 10.4110/in.2023.23.e28.

35. Chen, Z., Kong, X., Ma, Q., Chen, J., Zeng, Y., Liu, H., Wang, X. and Lu, S., 2024. The impact of *Mycobacterium tuberculosis* on the macrophage cholesterol metabolism pathway. Frontiers in Immunology, 15, p.1402024. doi: 10.3389/fimmu.2024.1402024.

36. Agarwal, P., Gordon, S. and Martinez, F.O., 2021. Foam cell macrophages in tuberculosis. Frontiers in Immunology, 12, p.775326. doi:10.3389/fimmu.2021.775326.

37. Ye, Y., Liu, J., Guo, Y., Gao, Y., Rao, J., Su, R., Zhang, L., Huang, Z., Luo, Q. and Li, J., 2022. PPARγ ameliorates *Mycobacterium tuberculosis* H37Ra-induced foamy macrophage formation via the ABCG1-dependent cholesterol efflux pathway in THP-1 macrophages. Frontiers in Microbiology, 13, p.829870. doi: 10.3389/fmicb.2022.829870.

